# Unpaired Data Empowers Association Tests

**DOI:** 10.1101/839159

**Authors:** Mingming Gong, Peng Liu, Frank C. Sciurba, Petar Stojanov, Dacheng Tao, George C. Tseng, Kun Zhang, Kayhan Batmanghelich

## Abstract

To achieve a holistic view of the underlying mechanisms of human diseases, the biomedical research community is moving toward harvesting retrospective data available in Electronic Healthcare Records (EHRs). The first step for causal understanding is to perform association tests between types of potentially high-dimensional biomedical data, such as genetic, blood biomarkers, and imaging data. To obtain a reasonable power, current methods require a substantial sample size of individuals with both data modalities. This prevents researchers from using much larger EHR samples that include individuals with at least one data type, limits the power of the association test, and may result in higher false discovery rate. We present a new method called the Semi-paired Association Test (SAT) that makes use of both paired and unpaired data. In contrast to classical approaches, incorporating unpaired data allows SAT to produce better control of false discovery and, under some conditions, improve the association test power. We study the properties of SAT theoretically and empirically, through simulations and application to real studies in the context of Chronic Obstructive Pulmonary Disease. Our method identifies an association between the high-dimensional characterization of Computed Tomography (CT) chest images and blood biomarkers as well as the expression of dozens of genes involved in the immune system.

---

**I**ncreasingly, data from Electronic Health Records (EHRs) in hospitals are becoming available to clinical researchers. Such massive collections contain various types of data from sources such as high-resolution imaging, genome sequencing, and physiological metrics. By studying such a large and diverse data, researchers can provide a holistic view of the underlying mechanisms of human diseases. For example, while a large proportion of human diseases are influenced by genetic variants [1–3], their mechanisms are not well understood [4–6]. To understand the mechanism, measuring other variables such as gene expression is required. Unfortunately, it is unlikely that all patients in the EHR have all measurement modalities. For example, due to the high cost of image acquisition and specimen maintenance, hospitals order those only when they are needed. Consequently, only the record of a few patients contains all data modalities, which reduces the power of association tests and increases the chance of false discovery.

Furthermore, a multidimensional phenotype can offer better sensitivity to the clinical and genetic underpinning of human diseases than a one-dimensional scalar phenotype [7–9]. For instance, high-dimensional features can be computed to summarize the folding pattern of the brain structure in Magnetic Resonance (MR) imaging [10], or the texture and distribution of the lung tissue destruction can be measured and summarized by Computed Tomography (CT) imaging [11, 12]. Those metrics are highly predictive of the diseases (e.g., Alzheimer’s disease [9, 13] and bipolar disorder [14, 15] for MR, and COPD [12] for CT). Relating that high-dimensional phenotype to genetic and genomic measurements provides more evidence for understanding the etiology of the disease.

In this paper, we present a new method to formally test the association between two types of potentially high-dimensional data that allows incorporating unpaired samples, i.e., samples with one data modality (see Fig. 1 for a schematic illustration). Our approach provides better control of falsepositive and, under some mild assumptions, increases the statistical power of the test. Unpaired data enables us to better estimate the null distribution, which results in more accurate control of the false positive rate. Furthermore, it allows us to leverage the underlying structure of the high-dimensional measurements, which consequently increases the power of the test. The proposed method, the Semi-paired Association Test (SAT), falls in the kernel machine framework [16–25]. More specifically, two variants of our method generalize the Variance Component Score Test (VCST) [16–20] and the Kernel Independent Test (KIT) [21–25] such that they can exploit unpaired data. The VCST is commonly used to test for heritability of a phenotype [16–20] and is implemented in popular software such as GCTA [26]. The KIT is widely used for statistical independence test in various scenarios [22, 25, 27, 28]. We provide a connection between those methods. Our proposed test makes unpaired data, previously wasted, available for discovering novel associations in massive uncontrolled datasets such as EHRs. Unearthing unnoticed associations assists in understanding the underlying mechanism of human diseases.

**Figure 1:**
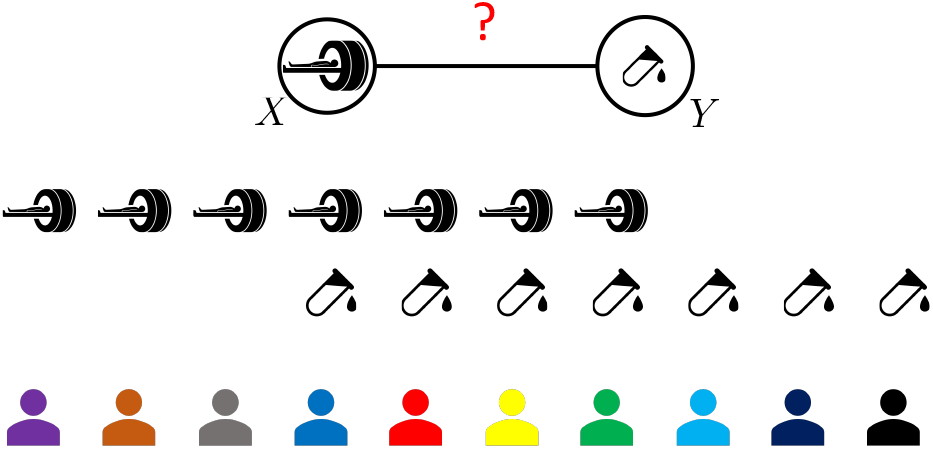
*X* and *Y* represent two modalities. Current approaches only use paired data 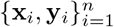. Assuming that the total number of samples of *X* (*M*) and *Y* (*N*) is more than the paired data, we aim to find out how the control of the false discovery and the power of association tests can be improved by the unpaired data 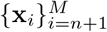 and 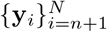.

This paper makes two contributions. First, it provides a statistically grounded method for the inclusion of unpaired data. The extensive simulation, as well as theoretical study, supports the hypothesis that the unpaired data is beneficial to control the false discovery and if the conditions are satisfied, can improve the power. Second, we apply our method to two different real studies. In the first experiment, we show that unpaired data can discover a new association between the high-dimensional radiographic measurements of Chronic Obstructive Pulmonary Disease (COPD) and peripheral blood biomarkers that play a role in the immune system. In this dataset, only a subset of the cohort has blood samples. In the second experiment, we apply our approach to genotype-phenotype data from the General Population Cohort from Uganda [29]. In this dataset, all subjects have genotype data but only one-fourth of them have phenotypes. Our method is able to find more heritable phenotypes.

## Results

### Theoretical Framework

We propose a method to test the association between two potentially high dimensional datasets. In addition to paired data, our method is able to exploit unpaired data, meaning the data from subjects that have only one modality. One can view the distance between the joint distribution and the product of the marginals as the strength of the association. To test association formally, the null distribution for the distance should be estimated. Our first theorem (Theorem 1) shows that the null distribution can be estimated more accurately using unpaired data. Hence, our method results in better control of the type I error. In addition to type I error, we show, in Theorem 2, that power can be improved if the data (of at least one modality) live on a lower dimensional space. Such an assumption is mostly the case for real data. For example, previous studies have shown that the space of Magnetic Resonance images of the brain can be modeled by a relatively low-dimensional manifold [30–32]. A similar assumption has been explored to model the low-dimensional space of gene expression for single-cell expression analysis [33, 34]. Unpaired data help us to estimate the low-dimensional space more accurately. Our new test statistic, which is a random variable calculated from sample data, exploits the low-dimensional assumption by taking advantage of the unpaired data.

Our method, the Semi-paired Association Test (SAT), generalizes two popular methods for association testing, the Variance Component Score Test (VCST) and the Kernel Independent Test (KIT), which are commonly used in statistical genetics to test the heritability of traits [17, 20] or gene-level associations [19, 28, 35]. More specifically, a variant of our method, SAT-fx, generalizes the VCST, which assumes that one of the modalities is not a random variable (i.e., fixed). For example in heritability analysis, the effect size is random but the genotype is given. The second variant, SAT-rx, generalizes KIT, which assumes that both modalities are random variables.

### Simulation Results

To evaluate our method’s improvement of type I and type II errors, we mimic the data missingness mechanism by conducting two levels of simulations:

i. We synthesize both *X* and *Y*. In this simulation, we evaluate both variants of our method, including SAT-fx and SAT-rx.
ii. Following the literature of population genetics in which testing for the heritability of traits is a topic of interest, we use genotype data as *X* and synthesize *Y*. We only evaluate SAT-fx because the genotype data is fixed.

In simulation (i), to generate *X*, we first generate *N* low-dimensional (dim=10) data points from a Gaussian distribution and then map them to high-dimensional *X* using a linear transformation plus independent Gaussian noise. To generate *Y*, we first generate low-dimensional data according to the variance components model (see Eq. 1 in the Method section) and then map them to high-dimensional *Y* using another linear transformation plus independent Gaussian noise.

In simulation (ii), we use the real genotype data from the COPDGene cohort as *X*. COPDGene is a multi-centered study of the genetic epidemiology of Chronic Obstructive Pulmonary Disease (COPD) that enrolled individuals aged 45-80 years with at least a 10 pack-year history of smoking [36]. We generate *Y* using the same procedure used in simulation (i).

In all the simulations, we create 1000 simulation replicates to evaluate the type I error rate and test power. Type I error rates and powers are calculated using the percentage of p-values smaller than a given significance level (*α* =0.05) under null models and alternative models, respectively. We set the heritability *h*^2^ = 0 for the evaluation of type I error rates and *h*^2^ =0.1 for the evaluation of power. To show the benefits of incorporating unpaired data, we compare the type I errors of the VCST/KIT as a baseline with two variants of both SAT-fx and SAT-rx: with and without Dimensionality Reduction (DR). VCST and KIT only use *n* paired data points, while SAT-fx and SAT-rx use *n* paired data points together with an additional *N* – *n* unpaired data points. For evaluation, we have access to the oracle where we can apply VCST and KIT using all *N* data points as paired, which is the best we can achieve. We set *n* = 100 for simulation (i) and *n* = 3000 for simulation (ii).

Fig. 2 and Fig. 3 report the type I error rates and power in simulation (i), respectively. Here we only show results for random *X*. The results for fixed *X* have similar trends and are available in Section S1 of the Supplementary Information. The results in Fig. 2 demonstrate that the type I error rates of our proposed method approach the predefined significance level (0.05) as we add more unpaired data. In addition, Fig. 3 shows that our method’s test power increases when adding unpaired data. Though our method has lower power than the oracle method which has access to all the paired data, it consistently outperforms the baseline KIT method that uses only paired data.

**Figure 2:**
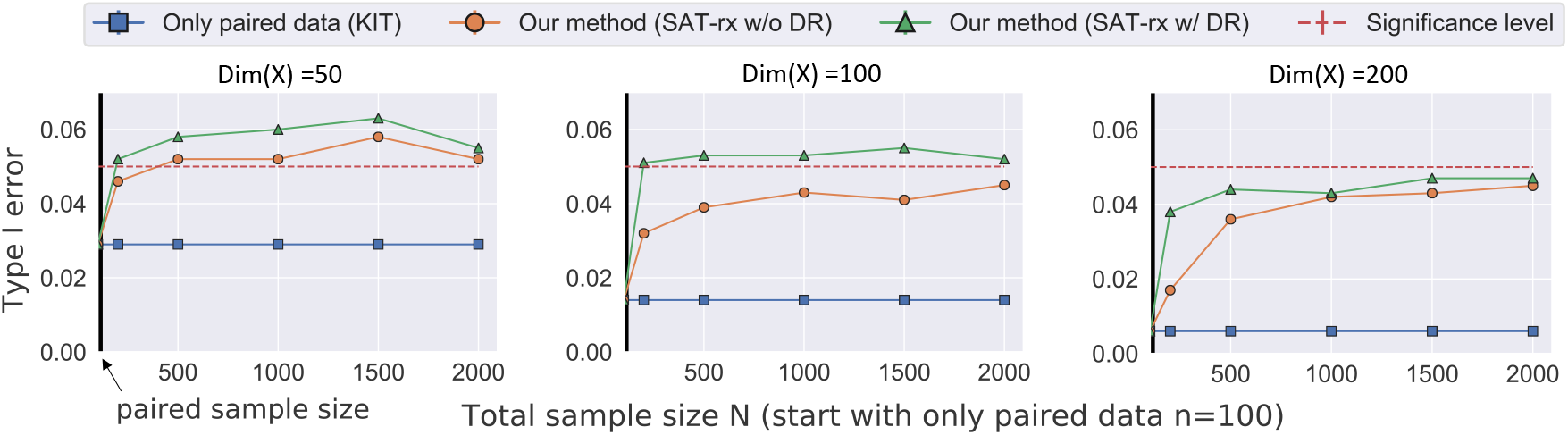
Evaluation of SAT-rx type I error rate control on the simulated data generated by procedure (i) in the random *X* setting. The blue line (KIT) is the result of using only paired data; hence it does not change with addition of unpaired data. KIT only uses the *n* = 100 paired data points. Our methods (green and orange) start with *n* pairs and gradually adds unpaired data to improve type I error control. False-positive rates for both variants of our method SAT-rx are well controlled around the nominal value (DR:Dimension Reduction).

**Figure 3:**
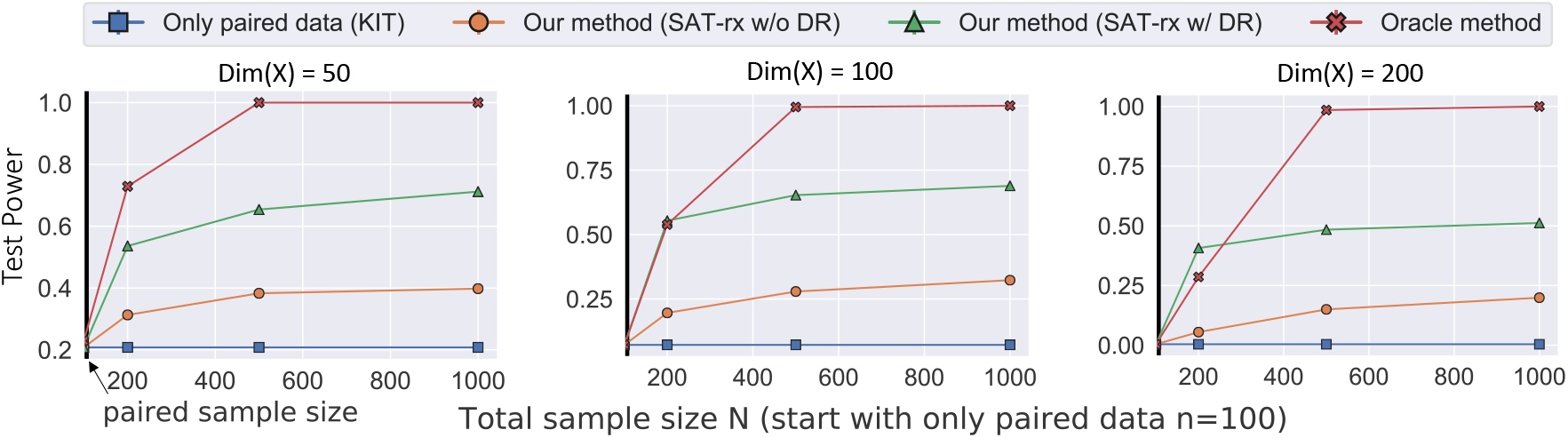
Evaluation of SAT-rx test power on the simulated data generated by procedure (i) in the random *X* setting (DR:Dimension Reduction). The results for heritability values *h*^2^ = 0.1 and dimensionality *dim*(*X*) = *dim*(*Y*) = 50, 100, 200 are shown. KIT only uses the *n* = 100 paired data points. Our methods start with *n* pairs and gradually add unpaired data to improve test power.

Fig. 4 and Fig. 5 report the type I error rates and powers of all the methods evaluated in simulation (ii). Again, we can see from Fig. 4 that the type I error rates of our proposed methods approach the significance level (0.05) as we add more unpaired data. However, because the dimensionality of the genotype is very high, the test is still very conservative even after adding unpaired data. Nevertheless, our method’s power exceeds that of VCST and increases as we add unpaired data.

**Figure 4:**
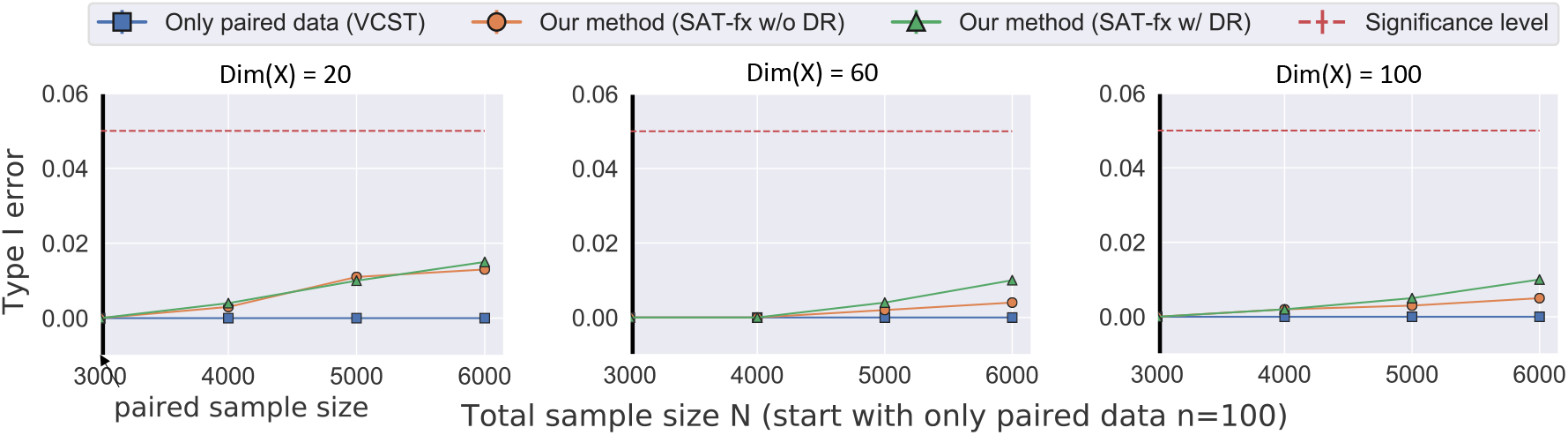
Evaluation of SAT-fx type I error rate control on the data generated in simulation (ii). VCST only uses the *n* = 3000 paired data points. Our method SAT-fx starts with *n* pairs and gradually adds unpaired data to improve type I error control.

**Figure 5:**
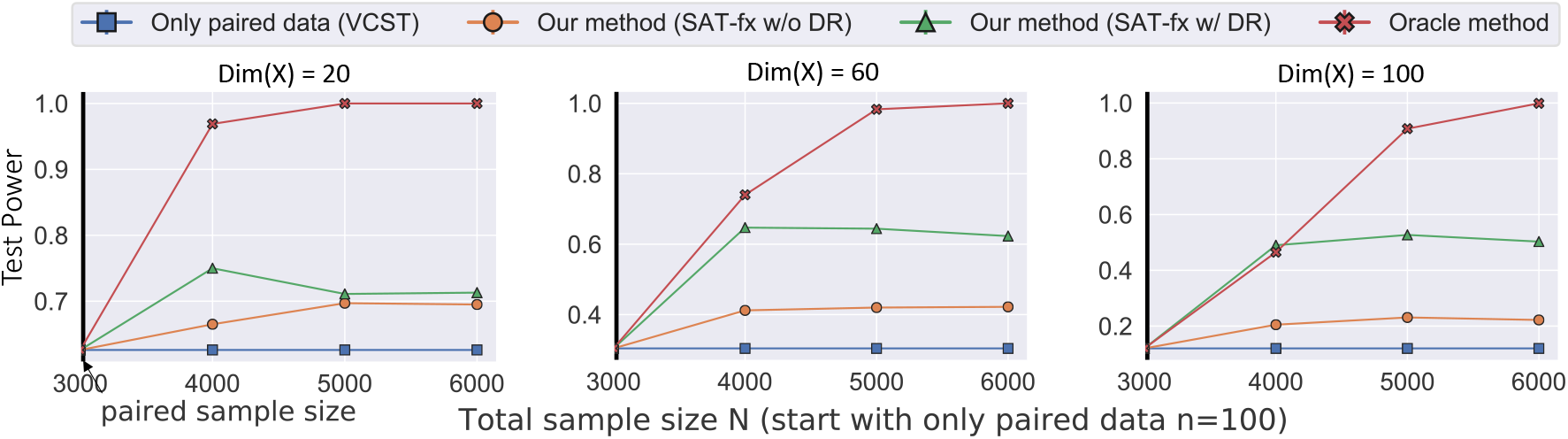
Evaluation of SAT-fx test power on the data generated in simulation (ii). VCST only uses the *n* = 3000 paired data points. Our method SAT-fx starts with *n* pairs and gradually adds unpaired data to improve test power.

### COPD: Imaging Data and Peripheral Blood Biomarkers

In this experiment, we investigate whether the high-dimensional radiographical measurement from Computed Tomography (CT) imaging is associated with peripheral blood biomarker signature of emphysema. COPD is a highly heterogeneous disease and involves many subprocesses, including emphysema [37]. CT imaging is increasingly used for emphysema diagnosis because it directly characterizes anatomical variation introduced by the disease [38]. Currently, Low Attenuation Area (LAA) is used to quantify the emphysema [39, 40]. However, LAA is based on a single intensity threshold value and cannot characterize variation in the texture of the lung parenchyma due to different disease subtypes [41]. Over the past year, researchers have proposed various generic and specific local image descriptors that extract higher order statistical features from CT images [11, 42, 43]. However, it is not clear whether such high-dimensional measurements are considered phenotypes, and whether the relationship to the causal biological processes is maintained.

We test the association between one of these multidimensional phenotypes and peripheral blood biomarkers. We use the method proposed by Schabdach *et al*. [11] that computes the similarity between 4629 patients and associates a 100-dimensional vector to each patient (see Supplementary Section S5 for details). Only 377 patients have both the blood biomarker and imaging data. We correct for the effects of covariates including age, sex, BMI (body mass index), and pack-year smoking history. Fig. 6 (a) reports the – log_10_(p-value) of different methods with respect to size of the unpaired imaging data. The results show that our method takes advantage of unpaired data and detects an association between high-dimensional imaging phenotypes and blood biomarkers that was not detected by the baseline method using only paired data.

**Figure 6:**
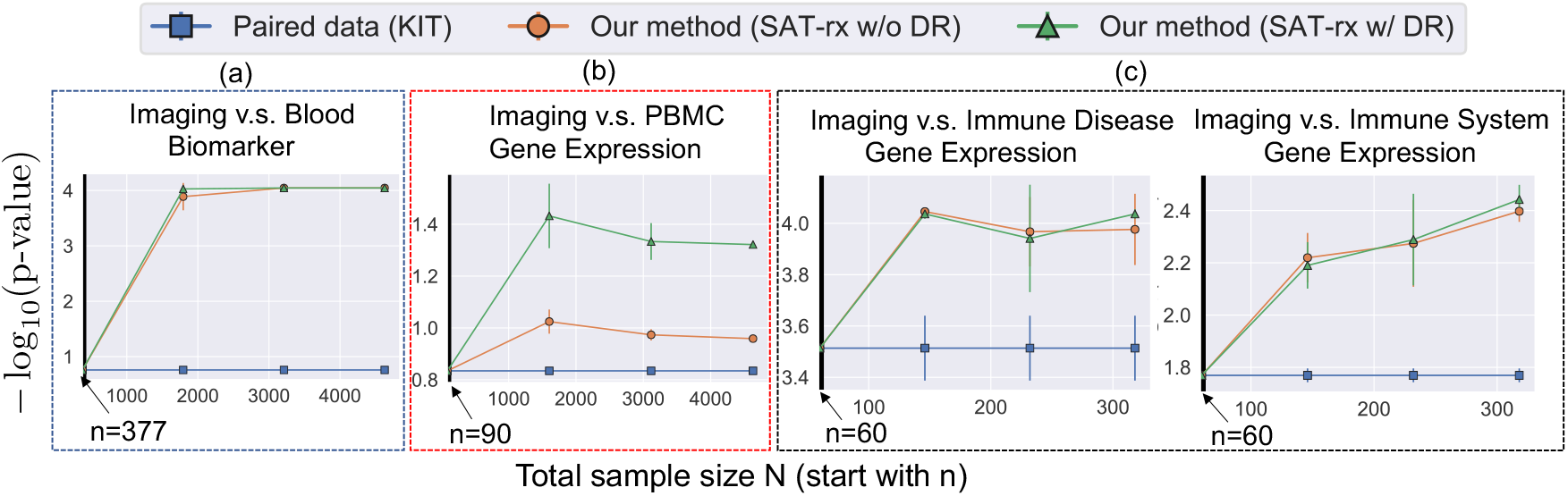
Experiments on three real imaging and genetics datasets. (a) Test an association between multidimensional imaging features and plasma biomarkers. (b) Test an association between imaging features and peripheral blood mononuclear cell gene expression data. (c) Test an association between imaging features and gene expression of genes in immune system pathway of the disease. In all the experiments, we start with *n* paired data points and show the behavior of our methods when adding unpaired data, with and without dimensionality reduction (DR).

### COPD: Imaging Data and Peripheral Blood Genes

Although smoking is a major risk factor for COPD, not all smokers develop debilitating disease, which suggests that COPD is a systemic disease and other factors might be involved in its development. Bahr *et al*. identified a set of genes whose expression is associated with two measurements used to diagnose COPD: percent predicted Forced Expiratory Volume in one second (FEV1) and the ratio of FEV1 to forced vital capacity (FEV1/FVC) [44]. These genes in Peripheral Blood Mononuclear Cells (PBMC) play a role in the immune system, inflammatory responses, and sphingolipid metabolism. Similar to the previous experiment, we investigate whether the multidimensional imaging phenotype is associated with systemic measurements. In this dataset, 90 subjects have both phenotype and gene expression measurements while more than 4539 subjects only have imaging phenotypes. We use the same covariates as the previous experiment. Fig. 6 (b) shows that our method exploits the unpaired data and results in lower p-values, suggesting an association between the imaging phenotypes and PBMC gene expression (p-value < 0.05) while the p-values of the baseline method using only paired data fails to pass the significance level.

### COPD: Imaging Data and Immune System Gene Expression

In this experiment, we apply our method again in the context of COPD but on a different dataset. We investigate the hypothesis that anatomical changes manifested on images are related to auto immune pathways. More specifically, we chose the “immune disease” and “immune system” gene pathways in the KEGG database [45]. We apply our method to imaging phenotypes and gene expression data containing 319 subjects from several sources (gene expression data from the GEO repository, imaging and clinical information from the Lung Genomics Research Consortium) [46]. Because only 60 patients have imaging phenotypes, we have a number of unpaired gene expression data. We compare our method with the baseline method that does not use the unpaired gene data and the results are shown in Fig. 6 (c). We can see that our method finds more significant associations as we add more unpaired data.

### Heritable Phenotype Discovery

In this section, we use the General Population Cohort (CPC), Uganda [29], to establish genotypephenotype associations in Genome-Wide Association Studies (GWAS), and show that our method can benefit from unpaired data.

GWAS have discovered many genetic risk variants of common diseases [2, 3]. Before performing GWAS, one should test the hypothesis that a given phenotype is “heritable” or not. Given the observation of a phenotype in a population of subjects, so-called narrow sense heritability is defined as an additive genetic portion of the phenotypic variance [47, 48]. A linear mixed model (LMM), which is a form of multivariate regression, is used to estimate the heritability (*h*^2^). Testing for the null hypothesis of 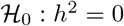 can be done using VCST and the power of the test is affected by the sample size.

We apply our method to study the heritability of a set of phenotypes from the General Population Cohort (GPC), Uganda. More specifically, it contains 37 phenotypes, including anthropometric indices, blood factors, glycemic control, blood pressure, lipid tests, and liver function tests (see the complete list of phenotypes in Supplementary section S6). Initially, 5000 individuals were genotyped using the Illumina HumanOmni 2.5M BeadChip array, out of which 4778 samples pass the quality control. We follow Heckerman *et al*. [49] exactly for quality control including the Hardy-Weinberg equilibrium (HWE) test, exclusion of Single Nucleotide Polymorphisms (SNPs) with low Minor Allele Frequency (MAF), and computation of the related matrix.

Among all the phenotypes, 18 phenotypes were measured for all the subjects, while the remaining 19 phenotypes were recorded for only 1423 subjects. Thus we conduct two sets of experiments for these two sets of phenotypes. For the 18 phenotypes measured for all individuals, we conduct experiments to mimic the random missingness of phenotypes. We subsample 3000 individuals as unpaired data and allocate the rest as paired data. We compare the p-values of the KIT as a baseline with two variants of SAT-rx, with and without dimensionality reduction. In this experiment, we are mimicking the missingness, hence we have access to the oracle, i.e., applying KIT using all data as paired, which is the best we can achieve and which we also compare with our method. The upper half of Table 1 reports the p-values generated by different methods for all evaluated phenotypes. We can see that the oracle produces much smaller p-values in general, while the baseline KIT method can hardly find significant associations. Our SAT-rx method clearly outperforms the KIT method and approaches the performance of the oracle on some phenotypes. Among the 18 phenotypes, our method finds 5 more heritable phenotypes than the baseline method at significance level 0.05.

**Table 1:** P-values on Uganda General Population Cohort. The newly found associations by our method at the significance level 0.05 are marked as bold. Since we mimic the missingness for phenotypes in the top part of the table, we are able to compare our performance with the oracle. In the bottom part of the table, a subset of the subjects has a missing phenotype; hence, the oracle columns are empty.

|  | KIT |  | SAT-rx (w/o DR) |  | SAT-rx |  | Oracle |  |
| --- | --- | --- | --- | --- | --- | --- | --- | --- |
|  | p-value | p-value (Bonf) | p-value | p-value (Bonf) | p-value | p-value (Bonf) | p-value | p-value (Bonf) |
| SBP | 0.293 | 1.000 | 0.224 | 1.000 | 0.128 | 0.897 | 0.010 | 0.195 |
| DBP | 0.091 | 0.928 | **0.031** | 0.537 | **7.25e-03** | 0.138 | < 1.00e-05 | < 1.90e-04 |
| BMI | 0.101 | 0.907 | **0.035** | 0.528 | **0.011** | 0.214 | < 1.00e-05 | < 1.90e-04 |
| WHR | 0.249 | 1.000 | 0.171 | 0.901 | 0.119 | 0.810 | 0.033 | 0.630 |
| Weight | 0.057 | 0.819 | **0.012** | 0.235 | **1.63e-03** | **0.031** | < 1.00e-05 | < 1.90e-04 |
| Height | 0.031 | 0.532 | 3.81e-03 | 0.072 | 1.74e-04 | **3.31e-03** | < 1.00e-05 | < 1.90e-04 |
| HC | 0.095 | 0.930 | **0.031** | 0.503 | **0.010** | 0.196 | < 1.00e-05 | < 1.90e-04 |
| WC | 0.127 | 0.928 | 0.057 | 0.662 | **0.022** | 0.345 | 1.20e-05 | 2.28e-04 |
| ALT | 0.204 | 0.920 | 0.172 | 0.646 | 0.106 | 0.617 | 1.76e-03 | 0.033 |
| Albumin | 0.117 | 0.983 | **0.046** | 0.593 | **0.024** | 0.395 | < 1.00e-05 | < 1.90e-04 |
| ALP | 0.442 | 1.000 | 0.419 | 1.000 | 0.318 | 1.000 | 0.261 | 1.000 |
| AST | 0.293 | 1.000 | 0.322 | 1.000 | 0.276 | 0.875 | 0.187 | 1.000 |
| Bilirubin | 0.046 | 0.629 | 0.027 | 0.390 | 8.43e-03 | 0.160 | < 1.00e-05 | < 1.90e-04 |
| Cholesterol | 0.024 | 0.448 | 2.25e-03 | **0.043** | 1.96e-04 | **3.72e-03** | < 1.00e-05 | < 1.90e-04 |
| GGT | 0.307 | 1.000 | 0.290 | 0.801 | 0.265 | 0.800 | 0.039 | 0.734 |
| HDL | 0.063 | 0.717 | **0.017** | 0.326 | **4.76e-03** | 0.090 | < 1.00e-05 | < 1.90e-04 |
| LDL | 0.012 | 0.222 | 6.10e-04 | **0.012** | 2.20e-05 | **4.18e-04** | < 1.00e-05 | < 1.90e-04 |
| Triglycerides | 0.242 | 1.000 | 0.164 | 1.000 | 0.126 | 0.880 | 6.76e-04 | 0.013 |
| HbA1c2 | 6.23e-03 | 0.118 | 3.66e-04 | **6.95e-03** | 1.80e-05 | **3.42e-04** | < 1.00e-05 | < 1.90e-04 |
| WBC | 6.95e-03 | 0.139 | < 1.00e-05 | < **2.00e-04** | < 1.00e-05 | < **2.00e-04** |  |  |
| RBC | 0.011 | 0.219 | 4.40e-05 | **8.80e-04** | < 1.00e-05 | < **2.00e-04** |  |  |
| Hemoglobin | 0.041 | 0.815 | 1.18e-03 | **2.36e-02** | 1.40e-05 | **2.80e-04** |  |  |
| HCT | 0.025 | 0.508 | 3.36e-04 | **6.72e-03** | < 1.00e-05 | < **2.00e-04** |  |  |
| MCV | 1.47e-03 | 0.029 | < 1.00e-05 | < 2.00e-04 | < 1.00e-05 | < 2.00e-04 |  |  |
| MCH | 2.50e-03 | 0.050 | < 1.00e-05 | < 2.00e-04 | < 1.00e-05 | < 2.00e-04 |  |  |
| MCHC | <1.00e-05 | <2.00e-04 | < 1.00e-05 | < 2.00e-04 | < 1.00e-05 | < 2.00e-04 |  |  |
| RDW | 6.70e-03 | 0.134 | < 1.00e-05 | < **2.00e-04** | < 1.00e-05 | < **2.00e-04** |  |  |
| PLT | 3.00e-03 | 0.060 | < 1.00e-05 | < **2.00e-04** | < 1.00e-05 | < **2.00e-04** |  |  |
| MPV | 1.00e-05 | 2.00e-04 | < 1.00e-05 | < 2.00e-04 | < 1.00e-05 | < 2.00e-04 |  |  |
| NEUPr | 0.015 | 0.304 | 7.80e-05 | **1.56e-03** | < 1.00e-05 | < **2.00e-04** |  |  |
| LYMPHPr | 3.30e-03 | 0.066 | < 1.00e-05 | < **2.00e-04** | < 1.00e-05 | < **2.00e-04** |  |  |
| MONOPr | 7.48e-03 | 0.150 | < 1.00e-05 | < **2.00e-04** | < 1.00e-05 | < **2.00e-04** |  |  |
| EOSPr | 1.13e-01 | 1.000 | **0.017** | 0.331 | **7.08e-04** | **0.014** |  |  |
| BASOPr | 9.60e-04 | 0.019 | < 1.00e-05 | < 2.00e-04 | < 1.00e-05 | < 2.00e-04 |  |  |
| LYMPH | 4.10e-04 | 0.008 | < 1.00e-05 | < 2.00e-04 | < 1.00e-05 | < 2.00e-04 |  |  |
| NEU | 0.062 | 1.000 | **3.31e-03** | 0.066 | **6.00e-05** | **1.20e-03** |  |  |
| MONO | 0.012 | 0.236 | 2.40e-05 | **4.80e-04** | < 1.00e-05 | < **2.00e-04** |  |  |
| EOS | 0.212 | 1.000 | 0.080 | 1.000 | **6.78e-03** | 0.136 |  |  |
| BASO | 1.20e-05 | 2.40e-04 | < 1.00e-05 | < 2.00e-04 | < 1.00e-05 | < 2.00e-04 |  |  |

For the other 19 phenotypes, 1415 individuals have both genotype and phenotype values, and the remaining individuals are considered unpaired (only genotype). We compare the p-values of the KIT as a baseline with two variants of SAT-rx, with and without dimensionality reduction. The lower half of Table 1 reports the p-values for all methods evaluated on these phenotypes. Among the 19 phenotypes, our method identifies 12 more heritable phenotypes than the baseline method at significance level 0.05.

## Discussion

Establishing the associations between various types of biomedical variables is essential for an understanding of disease mechanisms. When the biomedical variables are high dimensional, for example SNPs and imaging phenotypes, a huge sample size is required to guarantee enough statistical power. Also, a small sample size increases the chance of false discovery. However, due to the high cost of data collection, the available sample size is typically not sufficiently large, and the missingness in the data can cause a further reduction of sample size. To alleviate that problem, the biomedical research community is increasingly turning toward other sources (e.g., EHRs) where a massive amount of data is available. However, there is no guarantee that all subjects have all data modalities.

Here we address one type of missingness that frequently happens in the current association testing. When studying the relationships between two variables, one problem researchers usually encounter is that many data points have observations of only one variable (unpaired data), because the data for these variables may be collected initially for other purposes. We aim at being less “wasteful” by exploiting data points that have this type of missingness. Our method is based on a technical assumption that the data missing mechanism is independent of the association relationship, i.e., missing completely at random (MCAR). If this assumption is violated (e.g., we are only given biased paired data), our method cannot recover the original association. A future direction could be to extend our method to deal with more general missing mechanisms. Another primary assumption is that the distributions of the paired and unpaired data points are the same. For example, our method cannot be directly applied if gene expression data in the paired samples is collected by one platform while unpaired data is collected using a different platform. In this case, pre-processing is required to ensure that the platform bias is removed and the marginals of the paired and unpaired data are the same.

Although we showed the applicability of our method for univariate phenotypes (Table 1), the main focus of this paper is on multivariate phenotypes [10, 50]. Multivariate phenotypes provide more information than univariate ones, especially to study complicated phenomena such as the effect of cortical brain folding on the onset and progression of neurological diseases [9, 14, 51] and morphological traits in evolutionary biology [52]. Unpaired data enables us to discover the linear or non-linear relationship between variables in the multivariate phenotype.

Our method can take advantage of the unpaired data in two ways. First, our method improves the null distribution estimation by using unpaired data, which offers better control of the false discovery rate. As shown in Theorem 1, the estimation error of the null distribution depends on the total sample size, suggesting that incorporating unpaired data can readily improve the estimation of the null distribution. Second, we construct a new test statistic that explores the low-dimensional structure from unpaired data. We showed that under mild conditions, the new test has higher test power. It is worth noting that higher power can only be obtained if at least one type of data has a low-dimensional structure. Otherwise, our method can possibly have lower power than the baseline methods due to the removal of useful information. The low-dimensional assumption is a reasonable assumption for a multivariate phenotype. Measurements from highly structured data such as imaging are usually modeled as data points on a low dimensional manifold. For example, Gerber *et al*. [31] constructed a low dimensional brain manifold to study clinical variables such as age. Schabdach *et al*. [11] used manifold learning techniques to model information extracted from CT images of the lungs and showed that the low dimensional representation is highly correlated with the severity of the disease. Shi *et al*. [53] assumed gene expression data lay on a manifold and used a nonlinear dimensionality reduction method to capture biologically relevant structures in the data.

Our approach is closely related to the linear mixed effect model. The model is widely applied in statistical genetics to estimate the additive genetic effects of a univariate phenotype. Discovering a non-linear effect requires a much larger sample size, which might not be practical in terms of collecting enough data, at least when working with genotype data. However, there is no limitation in our approach’s ability to account for non-linear effects for other types of data. The linear effect assumption is equivalent to using linear kernels; different choices of the kernels (e.g., Radial Basis Function) can model non-linear effects. While the definition of heritability is well-defined for a univariate phenotype, it is less clear for a multivariate phenotype. We adopted the notion proposed by Ge *et al*. [10] (see Eq. 5). Their definition has several advantages. First, it is nicely connected to the mixed effect model and generalizes univariate heritability. Second, it allows us to incorporate unpaired data and derive the null distribution, which is required to compute the p-value efficiently, due to a mathematically appealing link with KIT and VCST. Deriving the null distribution for other definitions of multivariate heritability while incorporating the unpaired data (e.g., Zhou *et al*. [54]) is not as straightforward.

We conducted intensive simulation studies, evaluating various aspects of our proposed method. We generated synthetic data using the linear mixed effect model. To distinguish between SAT-fx and SAT-rx, we fix *X* in all the iterations for evaluating SAT-fx and randomly generate *X* in each replication for evaluating SAT-rx. Also, we provided a more practical simulation that uses real genotype data as *X* and only synthesizes *Y*. We set the heritability level (*h*^2^) to 10%, which is a modest heritability value. The higher values of *h*^2^ are less challenging than *h*^2^ =0.1 since the *X* and *Y* are more strongly related. The results in all the simulations demonstrate that our method better controls the type I error and has higher power than the baseline methods that ignore unpaired data. Fig. 4 suggests that when the dimensionality of *X* is large (e.g., *X* is genotype data) and the effect size is modest, our method controls the type I error conservatively and requires a larger sample size to reach the nominal level of type I error (i.e., 5%). Nevertheless, Fig. 5 shows that the gain in power using unpaired data is significant.

We also applied our method to the multivariate phenotype extracted from lung CT scans of patients with COPD. Two significant components of COPD are airway remodeling and alveolar destruction (emphysema) [55, 56]. Many pathogenic processes, such as chronic inflammation, contribute to the disease or cause anatomical variation [57, 58]. There is some evidence that the inflammatory process [58] and autoimmune response [59] are involved in emphysema. Researchers showed an association between various molecular signatures for COPD and emphysema in the peripheral blood mononuclear cells (PBMCs) [44, 60]. For example, Bowler *et al*. [60] investigated whether Interleukin-16, which is associated with autoimmune disease, is associated with COPD. Carolan *et al*. [61, 62] investigated the association of various blood biomarkers (e.g., adiponectin) with clinical and radiologic COPD phenotypes. However, those studies used FEV1 or LAA as surrogates for the disease severity, both of which are aggregate measures and cannot characterize a subprocess involved in the disease. For example, LAA is insufficient to distinguish emphysema visual subtypes because it merely counts the number of pixels in the lung region of CT images with intensity values lower than a single threshold. Since more sophisticated imaging descriptors (e.g., texture) are shown to be effective for emphysema sub-typing [63–65], we hypothesize that such descriptors are also associated with the systemic characterization of the disease such as PBMCs and gene expression. In this paper, we used a previously developed method [11] that uses image texture and intensity value and constructs a multivariate vector for each patient. The patient vector is shown to be more potent than LAA to characterize the disease severity [11]. We also construct a multivariate phenotype from rather traditional imaging measurements shown to be informative in previous studies [46]. We study the relationship between the imaging phenotype with the various blood measurements that are correlated with the activities of the immune system. Figure 6 reports that the classical approach in both experiments cannot detect the dependence while our method can. It is important to note that we test the dependency between a group of genes involved in the immune pathway, and our method does not identify specific causal genes contributing to the destruction of the tissue, which needs further investigation.

When computing the test statistic, our method adds additional computational load compared to the baselines VCST and KIT, due to the eigendecomposition of the kernel matrix on the paired and unpaired data. We generated samples from the null distribution to calculate p-values for both the baseline methods and our method, which is computationally expensive if a high precision p-value is required. Methods such as gamma approximation can be used to speed up the computation, which will of course introduce approximation errors.

## Method

In this section, we first give a brief review of the variance component score test (VCST) and the kernel independence test (KIT). We then discuss the connections between them and show that the differences between them lead to different ways to utilize unpaired data. Finally, we detail our SAT method by demonstrating how unpaired data can be incorporated to improve both VCST and KIT.

### Variance Component Score Test (VCST)

We start with the variance component model (a.k.a. the random effect model), which is widely used in statistical genetics for genetic association studies [10, 16, 26, 66]. We use the same nomenclature where 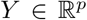 is a *p*-dimensional phenotype and 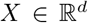 is genotype. However, our method is general and can be applied elsewhere. Given a paired sample containing *n* observations 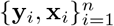, we consider the following multidimensional variance component model [10]:

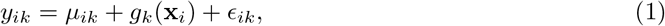

where *y_ik_* is the *k*-th element of **y**_*i*_, *g_k_* is a nonparametric function in a reproducing kernel Hilbert space (RKHS) associated with kernel *k*(**x**, **x**′) = 〈*ϕ*(**x**), *ϕ*(**x**′)〉, *μ_ik_* is the offset term, and *ϵ_ik_* is the error term. (1) can be rewritten in matrix form:

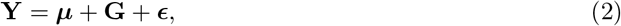

where 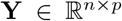 is the phenotypic matrix of the *n* observations (subjects) with *i*-th row 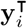, ***μ*** = (*μ*_1_, …, *μ_p_*) ⊗ **1**_*n*_ is a matrix of offsets (**1**_*n*_ is an *n* × 1 vector of ones), 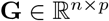 is the matrix of the aggregate genetic effects, and 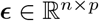 is a matrix of residual effects. We have the following distributional assumptions:

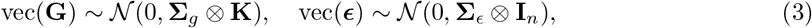

where vec(·) is the matrix vectorization operator that converts a matrix into a vector by stacking its columns, ⊗ is the Kronecker product of matrices, **I**_*n*_ denotes an *n* × *n* identity matrix, **Σ**_*g*_ is the genetic covariance matrix, **Σ**_*ϵ*_ is the residual covariance matrix, and **K** is the kernel matrix with *ij*-th element [**K**]_*ij*_ = *k*(**x**_*i*_, **x**_*j*_). For example, in the context of statistical genetics, **K** denotes identity-by-state (IBS) kernel [17,67,68], where [**K**]_*ij*_ represents the relatedness between individual *i* and *j*.

To test whether *Y* and *X* are associated (whether *Y* is heritable if *X* is the genotype), we can test the variance components as 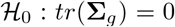 versus 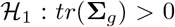 using the following score test statistic derived from model (1):

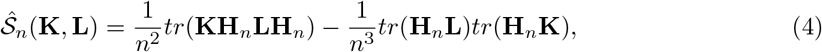

where *tr*(·) computes the trace of a matrix, 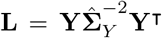 and 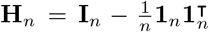, and 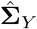 is the empirical covariance matrix of *Y*. The derivation details are provided in the Supplementary Information. The exact fraction of phenotype variability attributed to genetic variation is defined as heritability. There are various ways to define heritability for a multivariate phenotype (e.g., [10,54]). We adopt the definition by Ge *et al*. [10] that closely related to the VCST and subsumes the definition of the heritability for the univariate phenotype, which can be calculated as follows [10]:

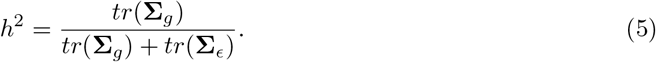

### Kernel Independence Test (KIT)

Kernel independence tests are a class of nonparametric methods which are also widely used for genetic association studies [22, 28]. Here we briefly review the Hilbert-Schmidt Independence Criterion (HSIC)-based independence test [22], which provides a general framework for many association tests [25]. Let 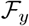 be a RKHS associated with the kernel function *l*(**y**, **y**′) = 〈*ψ*(**y**), *ψ*(**y**′)〉. HSIC tests 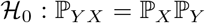 versus 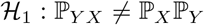 by testing 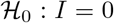 versus 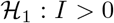, where *I* is defined as follows:

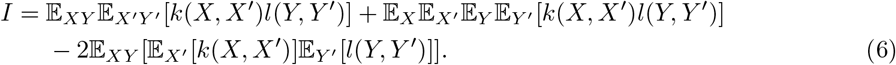

Given paired data of *n* subjects, an unbiased estimator of *I* is the following [69]:

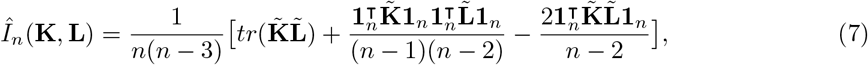

where 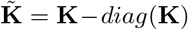 and similarly for 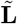 and **L**_*ij*_ = *l*(**y**_*i*_, **y**_*j*_). To test for statistical independence, one can use characteristic kernels, e.g., the radial basis function 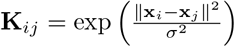, such that *I* can be zero only when *X* and *Y* are independent [70].

### Connections between VCST and KIT

Now we discuss the similarities and differences between VCST and KIT. Supplementary Table 2 displays the test statistics and null distributions of VCST and KIT.

#### Test statistic

It can be seen from Supplementary Table 2 that the biased statistics of VCST and KIT are identical to each other, if setting 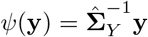. The unbiased test statistics of VCST and KIT differ. This is because VCST tests for random effects but assumes that the covariate inducing the random effect (*X*) and the corresponding kernel matrix (**K**) are fixed while KIT assumes *X* is random, leading to different ways to correct for the bias.

#### Null distribution

Let 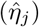 be the eigenvalues (empirical) of the covariance of *ϕ*(*X*) and let 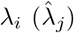 be the eigenvalues (empirical) of the covariance of *ψ*(*Y*). As shown in Table **??**, the null distributions for VCST and KIT have exactly the same forms, except that VCST uses 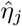 while KIT uses *η_j_*. This is also because of their respective fixed or random *X* assumptions. In practice, because λ_*i*_ and *η_j_* are both unknown, we need to replace them with 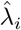 and 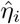. Therefore, the empirical null distributions of VCST and KIT are identical if only given *n* paired examples. However, they are inherently different because the null distribution of KIT is derived from asymptotic theory, while the null distribution of VCST is derived from the Gaussian error terms in the variance component model (2). This subtle difference is significant when using unpaired data, which is described as follows.

#### Unpaired data

The main difference between VCST and KIT is that *X* (**K**) is considered fixed or random respectively. When given unpaired data, VCST cannot make use of the unpaired data of *X* due to the fixed *X* assumption, while KIT can benefit from unpaired data of both *X* and *Y*. More specifically, unpaired data can only be used to improve the estimation of λ_*i*_ in VCST but they can be used to improve the estimation of both *η_j_* and λ_*i*_ in KIT.

### Semi-paired Association Test

In this section, we present our SAT method that incorporates unpaired data to improve test power. In addition to the *n* paired data, suppose we also have access to an unpaired sample 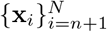 and an unpaired sample 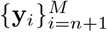. Without loss of generality, we assume *N* = *M* and replace *M* with *N* for notational simplicity. We will show two ways that unpaired data can improve the association test: 1) better control of type I error by improving the estimation of null distributions and 2) improved test power by devising a new test statistic under the intrinsic low-dimension assumption of high-dimensional data. We show how unpaired data are used for both VCST and KIT, resulting in two variants of our method, SAT-fx and SAT-rx.

#### Enhancing Type I Error Control

To calculate p-values, we need to estimate the parameters λ_*i*_ and *η_j_* in the null distributions from empirical data. Because λ_*i*_ and *η_j_* are the eigenvalues of the covariance of *ψ*(*Y*) and *ϕ*(*X*), respectively, the estimation does not require paired *Y* and *X* examples. Therefore, we can readily make use of unpaired data to obtain more accurate estimation of λ_*i*_ or *η_j_* involved in the null distribution.

For SAT-fx, we add unpaired *Y* data to estimate the covariance of *ψ*(*Y*) and its eigenvalues λ_*i*_ from both paired and unpaired data 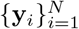, while *η_j_* should be estimated from only 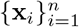 in the paired sample. For SAT-rx, we can further incorporate unpaired *X* data and use all the *X* data 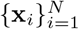 to estimate *η_j_*.

The following theorem shows that 1) the empirical null distribution convergences to the true (asymptotic) distribution and 2) the variance of the empirical null distribution converges to the variance of the true (asymptotic) null distribution with rate 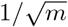, where *m* is the sample size of available data for estimating λ_*i*_ and *η_j_*.

##### Theorem 1 (Informal).

*Let* 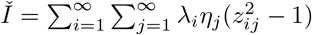 *and* 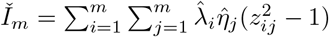.

1. *As m* → ∞, 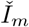 *converges in distribution to* 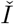.
2. *For all* 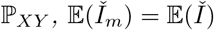 and 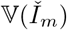 *converges in probability to* 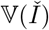 with rate 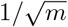.

The theorem is developed for SAT-rx and a similar theorem for SAT-fx can be considered as a special case of the above theorem. From the theorem, we can see that if only using paired data, *m* = *n*; if further using unpaired data, *m* = *N*. Because *N* > *n*, incorporating unpaired data to estimate λ_*i*_ and *η_j_* leads to lower estimation error and provides more accurate estimation of the null distribution. The proof details of Theorem 1 are given in Section 6 of the Supplementary Information.

#### Improving Test Power

Unpaired data contribute to a better estimation of the null distribution, resulting in better control of type I error. It can also improve test power. Specifically, if *X* or *Y* data (approximately) lie in a low-dimensional space, we show that unpaired data can be used to construct a new test statistic with improved test power. To devise the new test statistics, we first learn the low-dimensional space of *X* or *Y* by applying the kernel Principal Component Analysis (PCA) algorithm on both paired and unpaired data. Second, we project the paired data to the learned low-dimensional space and obtain the test statistics of our SAT-fx and SAT-rx by estimating the test statistics of VCST and KIT on the projected data. Due to the use of the kernel trick, calculating the test statistic of SAT-fx and SAT-rx requires only the kernel matrices **K**_*N*_ and **L**_*N*_ which are calculated on all the data, paired and unpaired.

In SAT-fx, because we do not consider *X* as random as does VCST, we can only incorporate unpaired *Y* data to learn the low-dimensional structure of *Y*. In SAT-rx, we further use unpaired data *X* to learn the low-dimensional space of *X*. The proposed new test statistics of SAT-fx and SAT-rx have the same form as that of VCST (4) and KIT(7), respectively. We only need to change the kernel matrices in the test statistics. Specifically, the new test statistic for SAT-fx is defined as 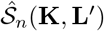, where

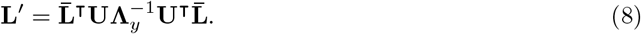

In **L**′, 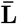 is the matrix comprised of the first *n* columns of **L**_*N*_, **U** = (**u**_1_, ⋯, **u**_*r_Y_*_) and 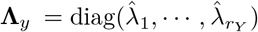 are the top *r_Y_* eigenvectors and eigenvalues of **L**_*N*_.

Similarly, the new test statistic of SAT-rx that considers *X* as random is 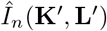, where

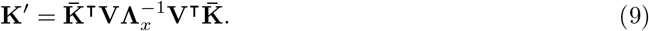

In 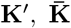 is the matrix composed of the first *n* columns of **K**_*N*_, **V** = (**v**_1_, ⋯, **v**_*r_X_*_) and 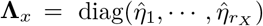 are the top *r_X_* eigenvectors and eigenvalues of **K**_*N*_. The asymptotic null distributions of the proposed 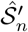 and 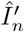 have the same forms as the null distributions of 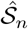 and 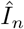, but using only the top eigenvalues 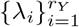 and 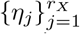, respectively. The derivation details are provided in Section 7 of the Supplementary Information.

The following theorem shows that the power of the new test statistic of SAT-rx is greater than the classical one that only uses paired data.

##### Theorem 2 (Informal).

*Assuming that data from X and Y lie in a low-dimensional manifold, the test power of the proposed SAT-rx is higher than that of the KIT method, which only uses paired data*.

SAT-fx follows similar properties as SAT-rx and can be considered as a special case of SAT-rx. The proof details of Theorem 2 are given in Section 8 of the Supplementary Information.

## Author Contributions

K.B. envisioned the project. M.G., K.B., and K.Z. developed the statistical method. M.G. implemented the method and performed the analyses. M.G. and K.B. wrote the paper. P.L. processed and performed analysis on the COPD imaging and immune system gene expression data. P.S. helped interpret the experimental results on medical data. D.T. provided assistance in theoretical analysis of the method. G.T. helped proofread biostatistical aspects of the paper.

## Supplementary Information

### S1. Simulation Results of SAT-fx

Fig. 1 and Fig. 2 report the type I error rates and power of SAT-fx, i.e., the method in the fixed X setting, in simulation (i), respectively.

**Figure 1:**
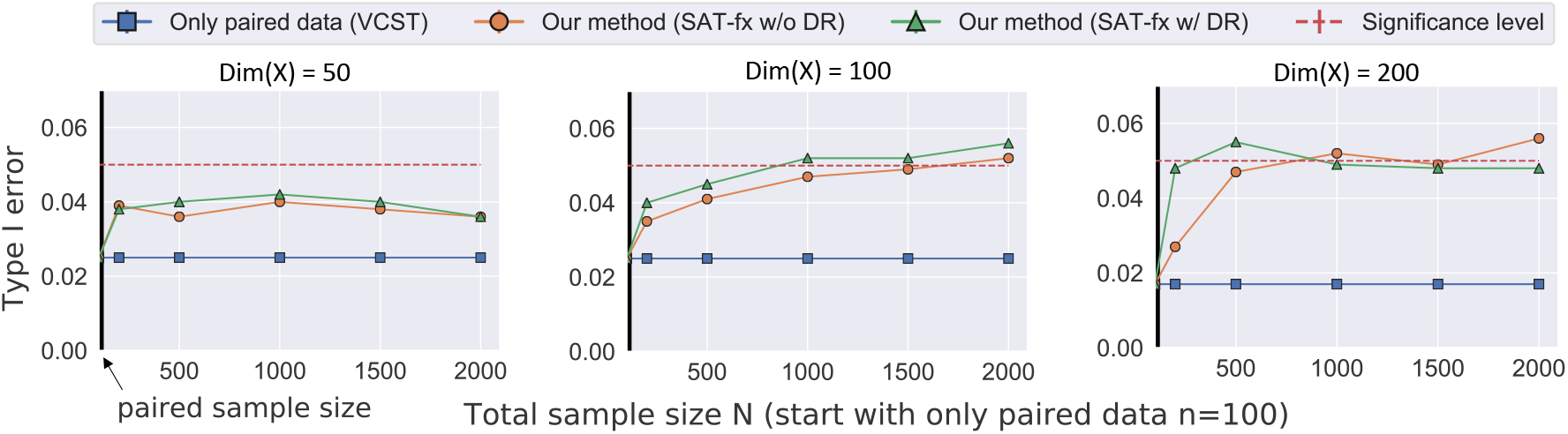
Evaluation of type I error rate control on the simulated data generated by simulation procedure (i) in the fixed *X* setting. The blue line (VCST) is the result of using only paired data; hence it does not change with addition of unpaired data. VCST only uses the *n* = 100 paired data points. Our methods (green and orange) start with *n* pairs and gradually adds unpaired data to improve type I error control. False-positive rates for both variants of our method SAT-fx are well controlled around the nominal value. (DR-Dimension Reduction)

### S2. Details about the Imaging Feature

We adopt the feature extraction strategy proposed by Schabdach *et al*. [1]. They proposed an efficient method that summarizes the CT images of patients to a low dimensional representation. They applied their method on a large cohort with COPD disease and showed that the low-dimensional representation is highly predictive of the disease severity. The general idea is to use a non-parametric method to first compute the similarity between pairs of patients, then construct the low dimensional representation which can be used to predict disease severity. In the following, we first explain the image pre-processing pipeline followed by the method used to compute the patient-patient similarity and the patient-level low-dimensional representation.

**Figure 2:**
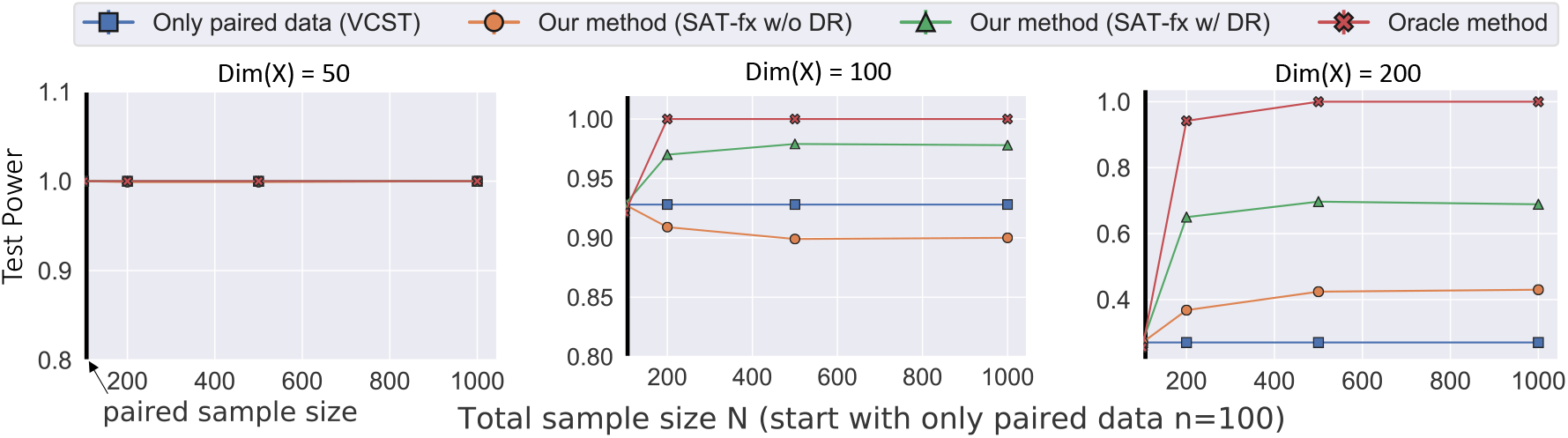
Evaluation of test power on the simulated data generated by procedure (i) in the fixed *X* setting. The results for heritability values *h*^2^ = 0.1 and dimensionality *dim*(*X*) = *dim*(*Y*) = 50, 100, 200 are shown. VCST only uses the *n* = 100 paired data points. Our methods start with *n* pairs and gradually add unpaired data to improve test power.

#### Pre-processing Pipeline

We apply the method to lung Computer Tomography (CT) images of 7,292 subjects from the COPDGene study [2]. The general pipeline is shown in Figure 3. First, we use SLIC [3] to over-segment the lung volume into the spatially homogeneous region, which is called super-voxelization. We extract texture and intensity features from each super-voxel. For the texture features, we use a method proposed by Liu *et al*. [4] called Spherical Histogram of the Gradients. It uses spherical harmonics to compute the histogram of gradients of pixels belonging to a super-voxel on a unit sphere. For the histogram features, we extract a 32-bin histogram of the intensity values of the pixels in a super-voxel. The intensity value of the CT images is shown to be highly informative for characterizing emphysema [5]. In summary, we model each patient as a bag-of-words [6] where the words are *d*-dimensional (*d* = 60) features extracted from super-voxels of lung CT image of a patient.

#### Computing Pairwise Similarities and Patient-Level Low Dimensional Representation

Let us denote *S_i_* = {*x*_*i*1_, ⋯, *x_iN_*} (*S_j_* = {*x*_*j*1_, ⋯, *x_jN_*}) be a set of all features from patient *i* (*j*) where *N* (*M*) represents total number of super-voxels of subjects *i* (*j*). We view *x_ik_* as random variable drawn from an unknown patient-specific probability density *p_i_* (*i.e*., *x_ik_* ~ *p_i_*). Schabdach *et al*. [1] proposed to use Kullback-Leibler divergence (KL) between *p_i_* and *p_j_* as a measure of pairwise *dissimilarity* between image data of patient *i* and *j*. The KL divergence is defined as

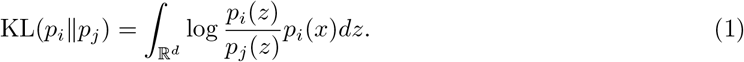

Instead of assuming an explicit parametrization, we follow Poczos *et al*. [7] that use a nonparametric estimator for KL divergence that is consistent and unbiased. Instead of global parametrization for *p_i_* and *p_j_*, they parametrize the probability densities *locally*. Let 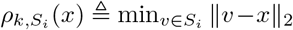 denote the 1 – *NN* distance from *x* in a set *S_i_*. Poczos *et al*. [7] proposed the following estimator for the KL,

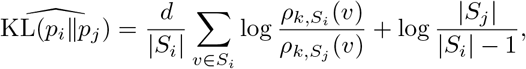

where |*S_i_*| and |*S_j_*| are the sizes of the sets *S_i_* and *S_j_* and *d* is the dimensionality of the input features.

**Figure 3:**
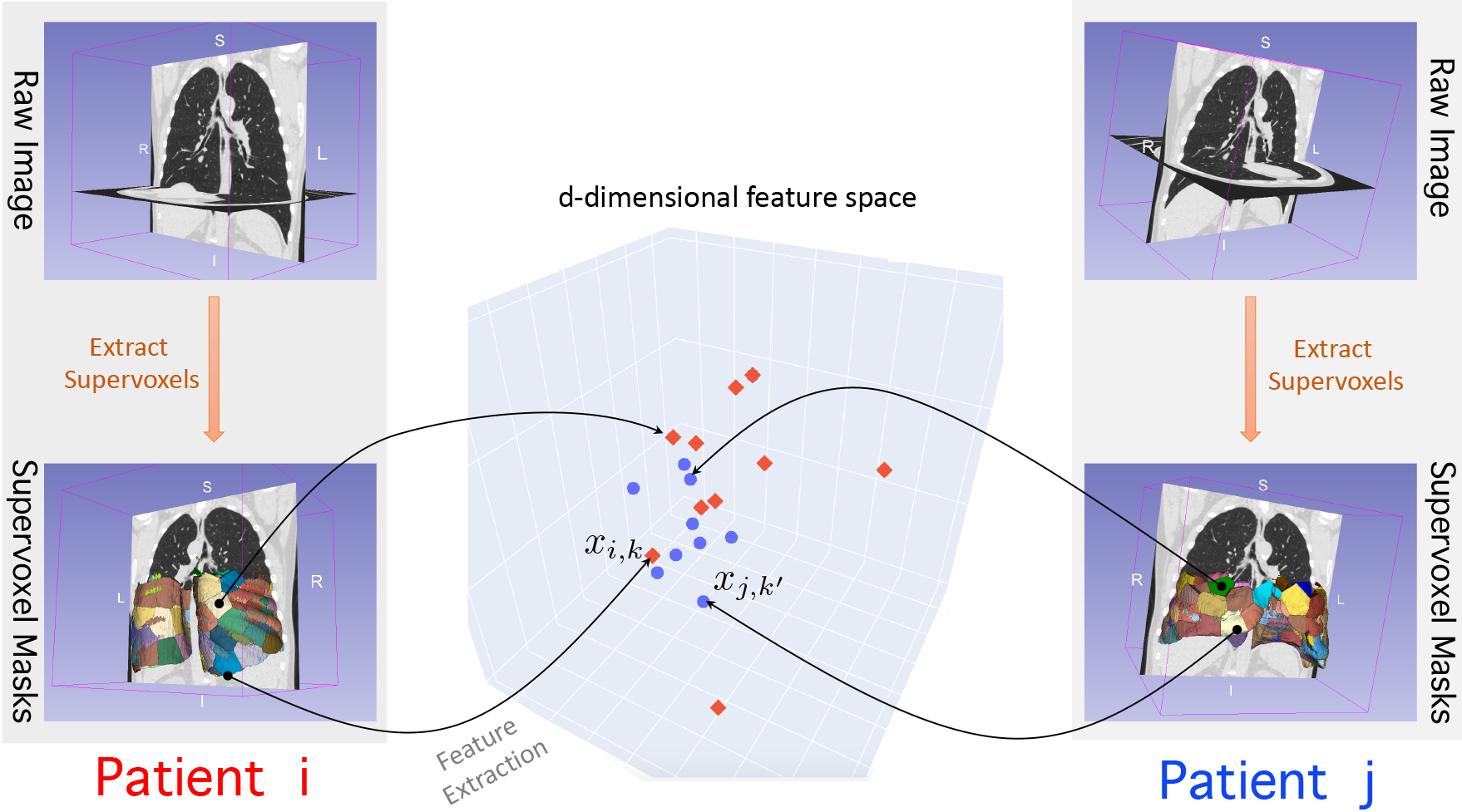
The schematic of feature extraction pipeline on lung imaging data. Each volumetric CT image is over-segmented (Extract Super-voxels). We extract a *d*-dimensional feature from each super-voxels. In this schematic, the blue circles (red rectangles) represent features extracted from super-voxels of the subject *j* (*i*).

The similarity kernel matrix is a Positive Semi-Definite (PSD) matrix. However, the KL divergence is neither symmetric nor a proper metric. First, we compute a matrix where the entry in row *i* and column *j* is

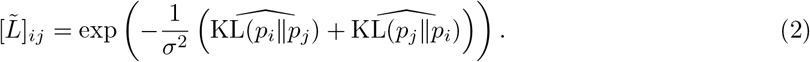

The *σ* is set to median of KL divergences (so-called median trick [8, 9]). Then, we project this matrix on the PSD cone to construct the kernel, 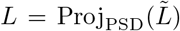, where Proj_PSD_ computes the Singular Value Decomposition of the input matrix and set the negative singular values to zero. Finally, we use the pairwise similarity kernel and apply Locally Linear Embedding (LLE) [10] to reduce the dimensionality and compute the patient-level feature representation.

### S3. Details about the Phenotypes in the Uganda Cohort

Table 1 explains the detailed information about each phenotye in the Uganda Cohort.

**Table 1:**
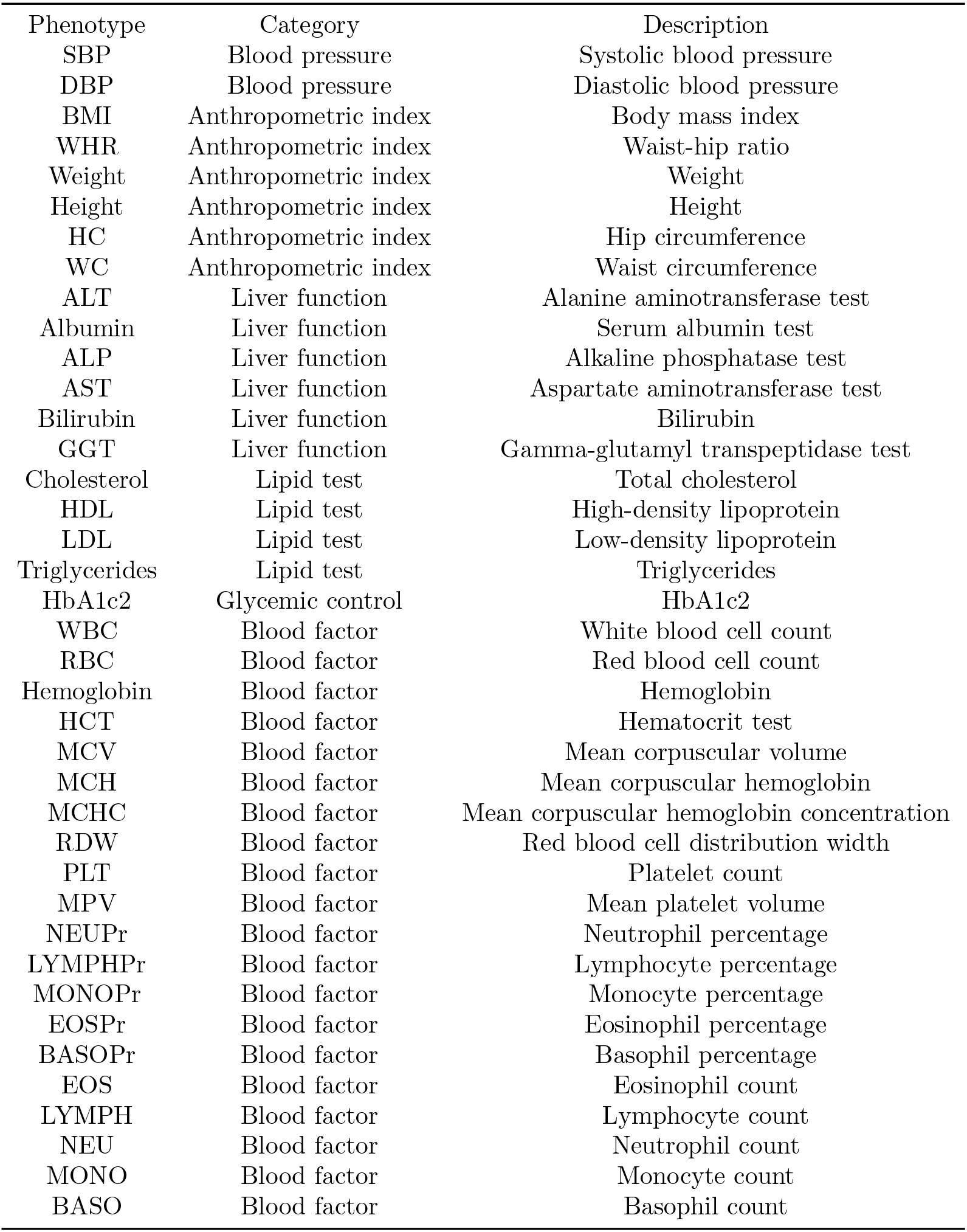
A description of the phenotypes measured in the Uganda cohort.

### S4. Comparison of VCST and KIT

Table 2 compares the test statistics and null distributions of VCST and KIT.

**Table 2:**
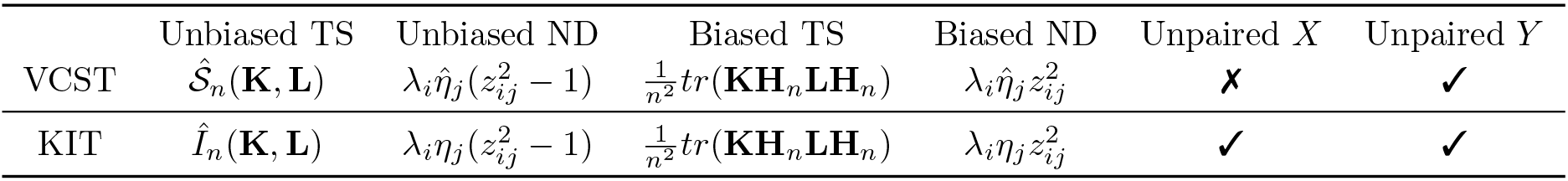
Comparison of VCST and KIT. TS: Test Statistic. ND: Null distribution.

### S5. Derivation of VCST Test Statistic 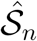 and the Null Distribution

Let us define 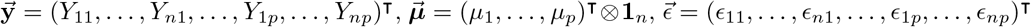, and 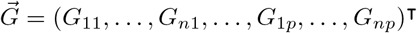, we can write the multivariate variance component model (Eq. 4 in the main text) as

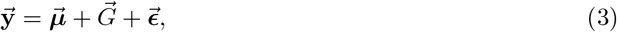

where 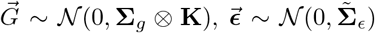, and 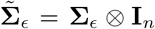. Therefore, we have 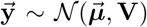, where 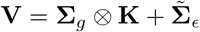. The corresponding restricted maximum likelihood (REML) is

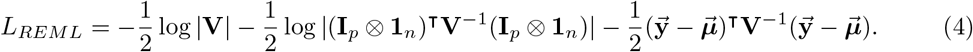

According to previous studies [11, 12], the score statistic evaluated at *H*_0_ can be defined as

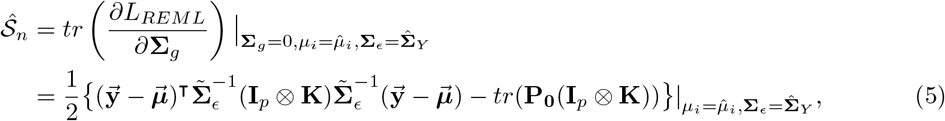

where 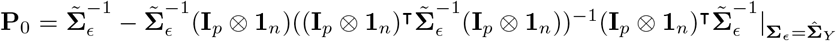. Equivalently, 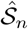 can be reformulated as follows

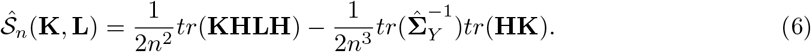

where 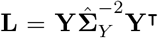. To derive the null distribution of 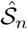, we reformulate the score statistic as 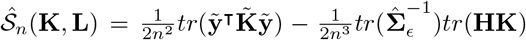, where 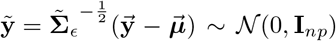 and 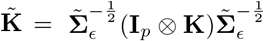. Let 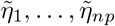 be the eigenvalues of 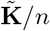. The eigenvalues can be calculated from the eigenvalues of **K** and 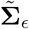 by 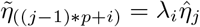, where λ_*i*_ are the eigenvalues of 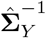. We then have 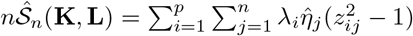.

### S6. Proof of Theorem 1

We first give a more formal statement of Theorem 1 in our main paper.

#### Theorem 1 (Formal).

Let 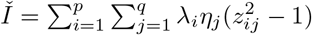 and 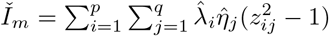.

1. Assume 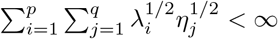. Then, as 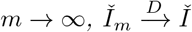.
2. 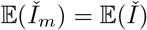. *For m* > 1 *and all δ* > 0, *with probability* 1 – *δ, for all* ℙ_*XY*_,

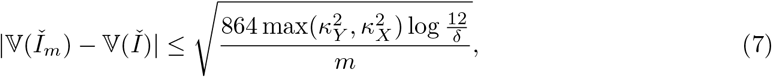

*where η_j_ be the eigenvalues of C_X_* (*covariance of ϕ*(*X*)*), 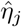 be the eigenvalues of Ĉ_X_ (empirical covariance of ϕ*(*X*)), λ*_i_ be the eigenvalues of the C_Y_ (covariance of ψ*(*Y*)*), and 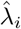 be the eigenvalues of Ĉ_Y_ (empirical covariance of ψ*(*Y*)*), respectively, in descending order. κ_Y_,κ_X_ are constants*.

*Proof*. (1) The proof of (1) in our Theorem 1 can be obtained by extending the proof of Theorem 1 in [13]. To prove 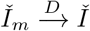, it suffices to prove

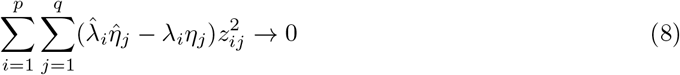

and

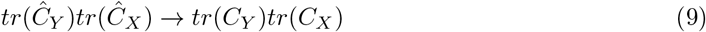

in probability as *m* → ∞. The convergence of the covariance trace operator has been proved in [13], i.e., 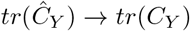 and 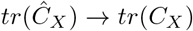. According to the continuous mapping theorem [14], we can immediately obtain (9). To prove (8), we can first get an upper bound

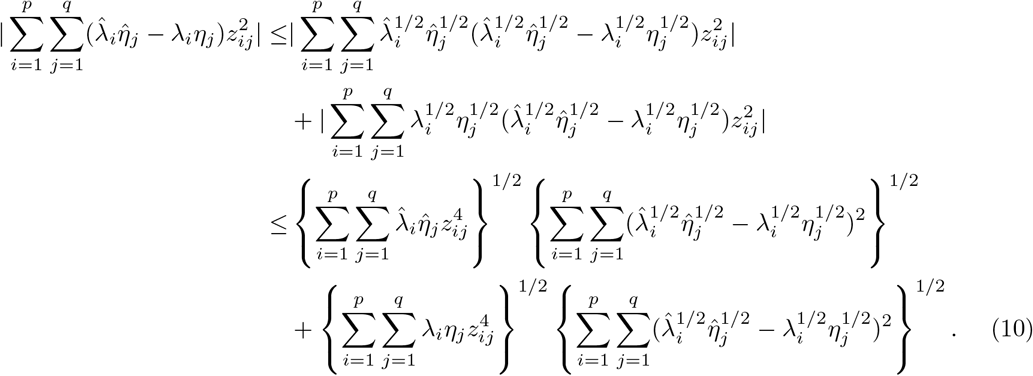

According to Chebyshev’s inequality, 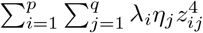 is of *O_p_*(1). Since 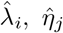, and *z_ij_* are independent, 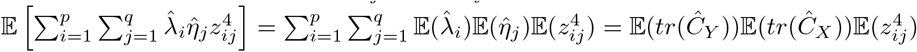. Because 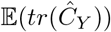 and 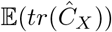 are bounded, we also have that 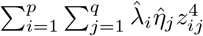 is of *O_p_*(1) according to Chebyshev’s inequality. The proof is complete if we show

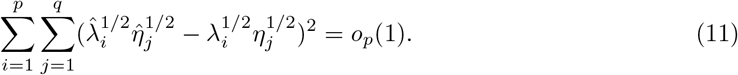

From 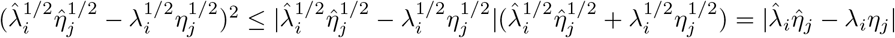, we have

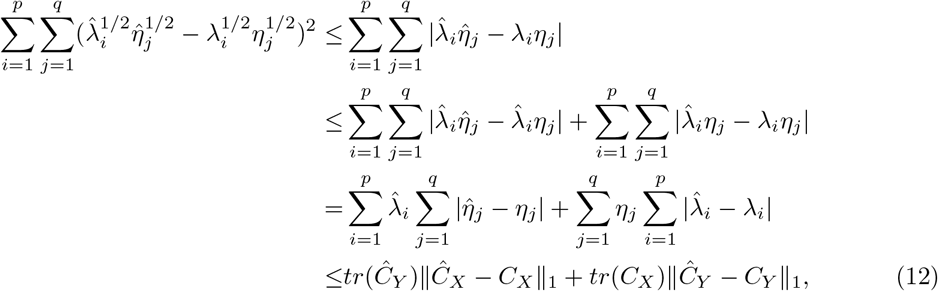

where ∥ · ∥_1_ is the trace norm. The last inequality makes use of generalized Hoffmann-Wielandt inequality. According to [15, Proposition 12], 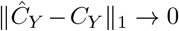 and 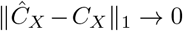 in probability, then the proof completes.

(2) To prove (2) in our Theorem 1, we first introduce the following Theorem on deviation bounds for U-statistics [16], which was obtained by applying a bound from [17, p. 25].

#### Theorem S1. (Deviation bound for U-statistics).

*A one-sample U-statistics is defined as the random variable*:

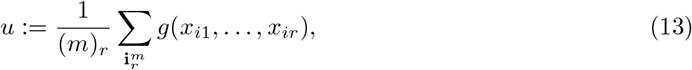

*where g is the kernel of the U-statistic. If a* ≤ *g* ≤ *b, then for all t* > 0 *the following bound holds:*

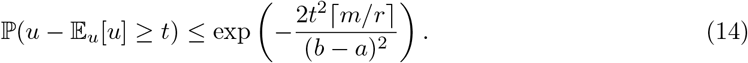

Now we are ready to prove the following Lemma.

#### Lemma S1.

*Given a random variable X with covariance in RHKS* 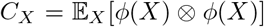 *and its empirical estimation* 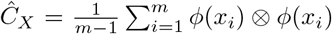 *from a sample S_x_* = {*x*_1_, …, *x_m_*}. *For all ϵ* > 0, *we have*

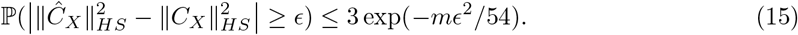

*Proof*. We first write 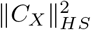 in terms of kernels:

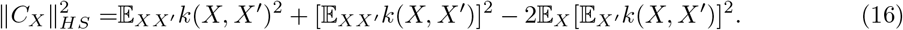

Similarly, we can also write 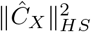 in terms of kernels:

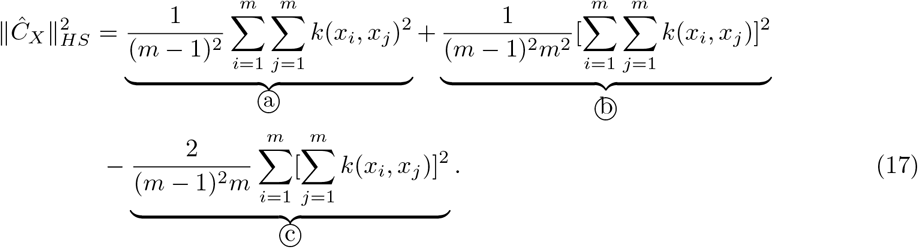

We first expand 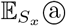 into

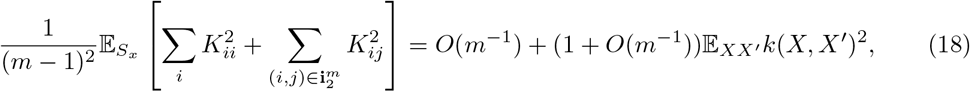

expand 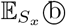 into

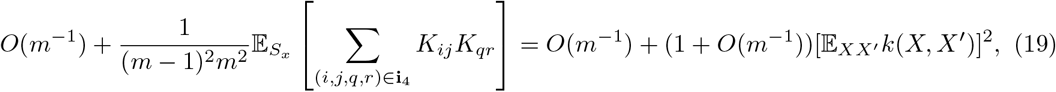

and expand 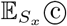 into

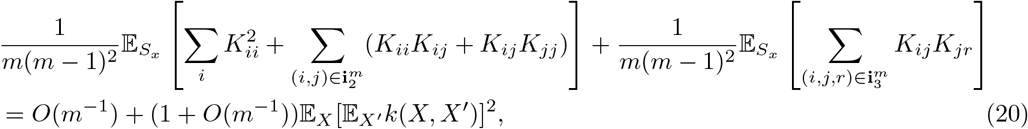

where 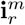 is the set of all r-tuples drawn without replacement from {1,…,m} and 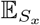 denotes the expectation w.r.t. *m* independent copies *x_i_* drawn from ℙ_*X*_. By omitting the terms that decay as *O*(*m*^−1^) or faster, we have

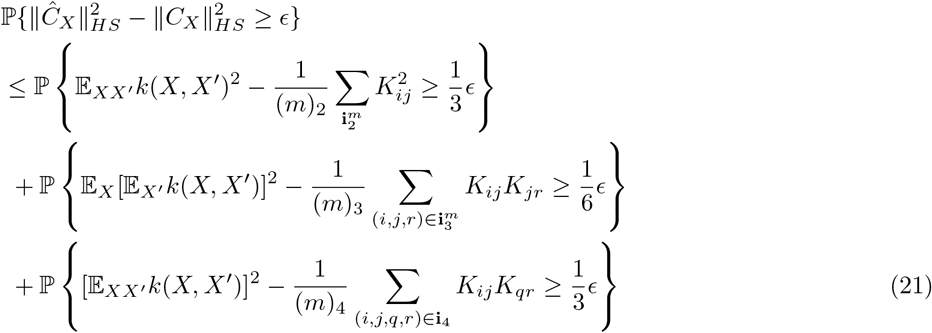

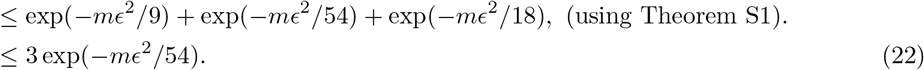

Now we are ready to prove (2) in our Theorem 1. From the definition of 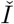 and 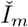, we have 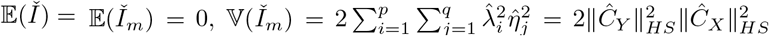 and 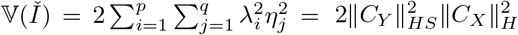. Then we have

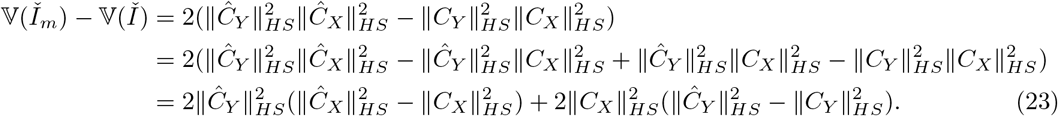

Thus,

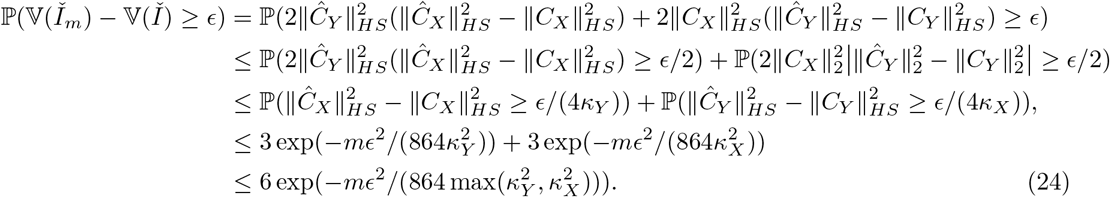

where 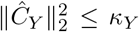 and 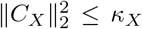. By setting 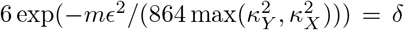, we can solve for *ϵ*:

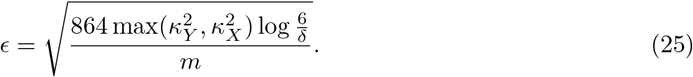

Therefore, we have that with probability at least 1 – *δ*,

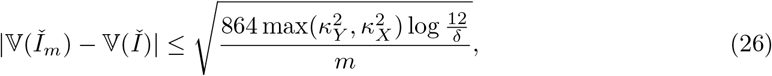

then the proof completes.

### S7. Derivation of the Null Distributions of Our SAT Test Statistics

Recall that our method SAT has two variants SAT-fx and SAT-rx, which are the extensions of VCST and KIT, respectively. Our SAT method basically project the original data into a subspace learned from unpaired data, and then plug in the projected data into the original VCST or KIT test statistics. The null distributions of our SAT-fx or SAT-rx test statistics follow the same forms as VCST or KIT, and differ in the number of *χ*^2^ terms in the summation. In the following, we will derive the null distributions of SAT-fx and SAT-rx test statistics separately.

#### SAT-fx

Let *U* be a the projection matrix containing first *r_Y_* columns of the eigenvector matrix of 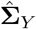, in which the eigenvectors are sorted according to their corresponding eigenvalues in a descent order. Let 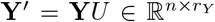. Here we define 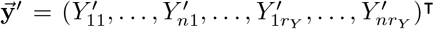, 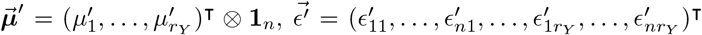. The covariance of *Y^′^* is defined as 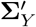 and the covariance of 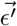 is defined as 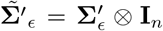. According to the derivation of the null distribution of 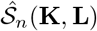 in Section S1, we reformulate the score statistic of our SAT-fx as 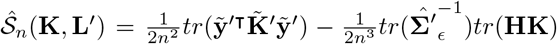, where 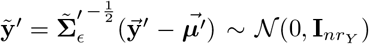 and 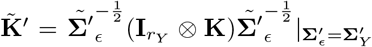. Let 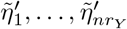 be the eigenvalues of 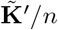. The eigenvalues can be calculated from the eigenvalues of **K** and 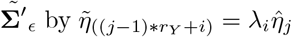, where λ_1_, …, λ_*r_X_*_ are the smallest *r_Y_* eigenvalues of 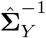. We then have 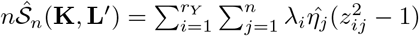.

#### SAT-rx

According to Mercer’s theorem [18], the kernel functions 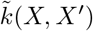 and 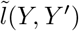 can be represented using eigenfunctions and eigenvalues defined in connection with them:

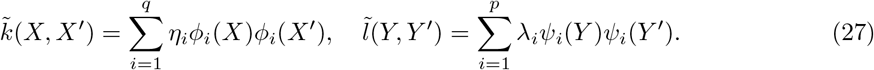

To derive the asymptotic null distribution of 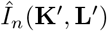, it suffices to derive the null distribution of 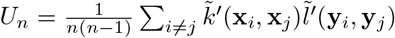, where

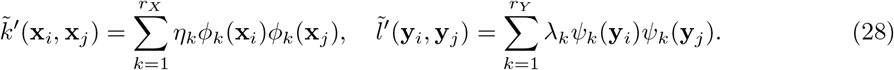

Using (28), we can rewrite *U_n_* as

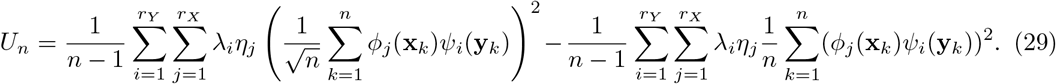

Let 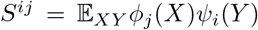 and 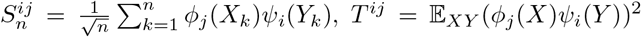, and 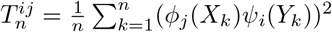, where 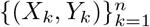 are *i.i.d*. variables with the same distribution as (*X, Y*). Under the null hypothesis, *S^ij^* = 0, the expectation of 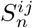 is

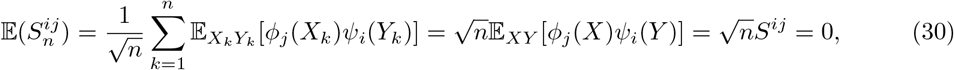

and the variance of 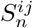 is

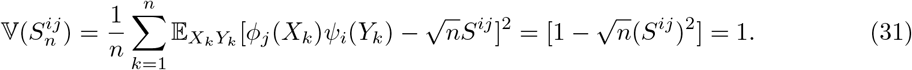

Similarly, the expectation of 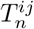 is

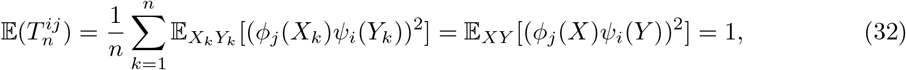

and the variance of 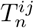 is

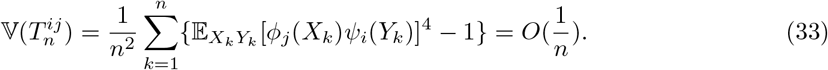

Thus, we have 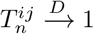 and 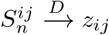, where *z_ij_* are standard normal variables. Therefore,

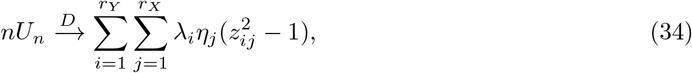

so does 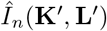.

### S8. Proof of Theorem 2

Here we give a formal version of Theorem 2 in our main paper.

#### Theorem 2 (Formal).

*We assume the following data generating process for X and Y:*

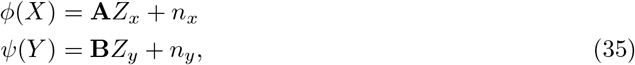

*where* 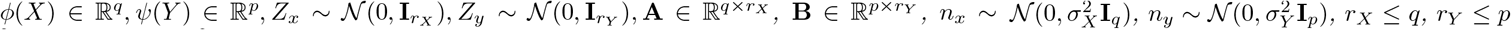, *and n_x_ is independent of n_y_. Under the alternative hypothesis, we have*

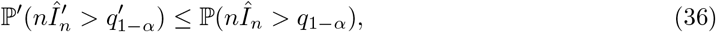

*where q*_1–*α*_ *and* 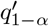 *are the* 1 – *α quantiles for the null distributions of* 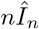 *and* 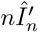, *respectively*.

*Proof*. We first give the results on the asymptotic distribution of 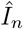 in the following lemma, which can be obtained by applying [19, Theorem 5.5.1 (A)].

#### Lemma S2.

*Let* 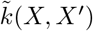 *and* 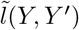 *be the centered kernel functions of k*(*X*, *X*′) *and l*(*Y*,*Y*′), *respectively. Assume that* 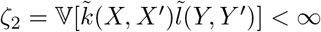 *and* 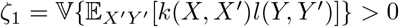. *Under the alternative hypothesis (I* > 0), *we have* 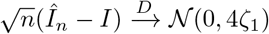.

Now we derive the asymptotic distribution of 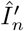 under the alternative hypothesis. Our method can be considered as the original KIT method with new kernel functions 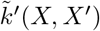 and 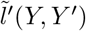 defined on the dimension-reduced inputs. Therefore, we have 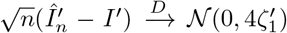, where 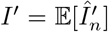. According to Mercer’s theorem [18], the kernel functions 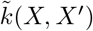 and 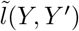 can be represented using eigenfunctions and eigenvalues defined in connection with them:

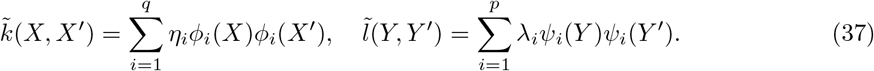

Similarly, the new kernels in the test statistic 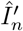 of our SAT-rx method can be represented as

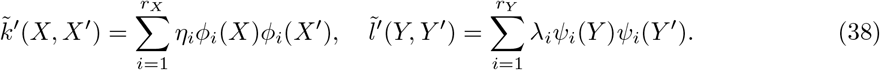

By using the representations of the kernels, we can express *I* as

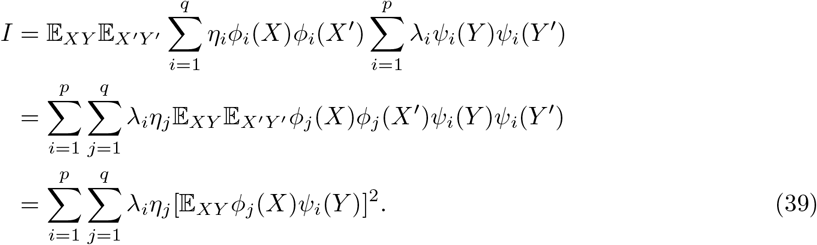

Similarly, 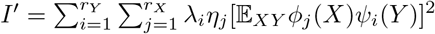. According to the connection between principal component analysis and the factor analysis model [20,21], the first *r_X_* principal component solutions correspond to the subspace spanned by **A** and the eigenfunctions of *C_X_* associated with the *q* – *r_X_* smallest eigenvalues map the input *X* to the noise term *n_x_*. Similarly, the eigenfunctions of *C_Y_* associated with the *p* – *r_Y_* smallest eigenvalues map the input *Y* to the noise term *n_y_*. Because *n_x_* and *n_y_* are independent, we have 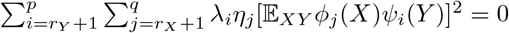, which implies *I* = *I*′.

By using the same representation, we can derive the relation between *ζ*_1_ and 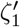. Using (37), *ζ*_1_ can be expanded as

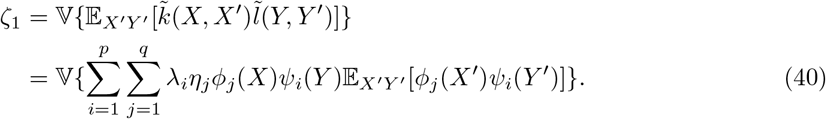

Similarly, using (38), we have

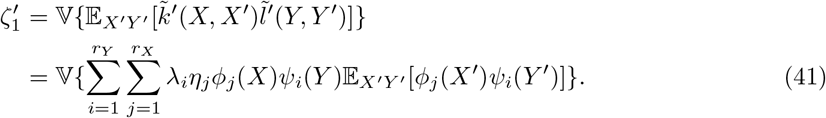

Because *r_X_* ≤ *q*, *r_Y_* ≤ *p*, we have 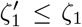. The power of the baseline KIT and our SAT methods at the significance level *α* can be calculated as 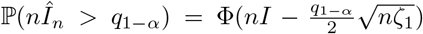 and 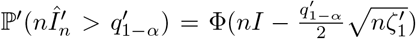, respectively, where Φ(·) is the CDF of a standard normal distribution, *q*_1–*α*_ is the 1–*α* quantile of 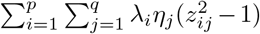, and 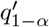 is the 1–*α* quantile of 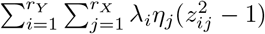. Because *r_X_* ≤ *q* and *r_Y_* ≤ *p*, we have 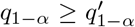, and further because of 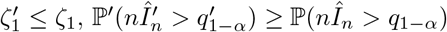. The proof completes.

